# Quantitative studies of an RNA duplex electrostatics by ion counting

**DOI:** 10.1101/645697

**Authors:** Magdalena Gebala, Daniel Herschlag

## Abstract

Ribonucleic acids are one of the most charged polyelectrolytes in nature, and understanding of their electrostatics is fundamental to their structure and biological functions. An effective way to characterize the electrostatic field generated by nucleic acids is to quantify interactions between nucleic acids and ions that surround the molecules. These ions form a loosely associated cloud referred as an ion atmosphere. While theoretical and computational studies can describe the ion atmosphere around RNAs, benchmarks are needed to guide the development of these approaches and experiments to-date that read out RNA-ion interaction are limited. Here we present ion counting studies to quantify the number of ions surrounding well-defined model systems of 24-bp RNA and DNA duplexes. We observe that the RNA duplex attracts more cations and expels fewer anions compared to the DNA duplex and the RNA duplex interacts significantly more strongly with the divalent cation Mg^2+^. These experimental results strongly suggest that the RNA duplex generates a stronger electrostatic field than DNA, as is predicted based on the structural differences between their helices. Theoretical calculations using non-linear Poisson-Boltzmann equation give excellent agreement with experiment for monovalent ions but underestimate Mg^2+^-DNA and Mg^2+^-RNA interactions by 20%. These studies provide needed stringent benchmarks to use against other all-atom theoretical models of RNA-ion interactions, interactions that likely must be well accounted for structurally, dynamically, and energetically to confidently model RNA structure, interactions, and function.

## INTRODUCTION

Ribonucleic acid (RNA) performs numerous functions in cells, including the storage and transmittal of genetic information, the regulation of gene expression, and catalysis, and each of these functions is fundamentally affected by the RNA’s high negative charge (1, 2). RNA carries one negative charged phosphoryl group per residue; hence, biologically-relevant RNAs that comprise of hundreds of nucleotides accumulate large and idiosyncratic charge densities.

Bringing RNA charges in close proximity during folding and function requires overcoming an enormous electrostatic energy barrier (3, 4). Ions, specifically cations, can reduce the electrostatic repulsion, which is referred to as screening (5-8) and, as importantly, mitigate electrostatic attraction with oppositely charged molecules, such as RNA binding proteins and aminoglycosides (9-12). Charge screening by ions is greatly affected by the charge of the cation in addition to its bulk concentration (13, 14), effects that are manifest in the folding of RNAs upon addition of millimolar Mg^2+^ in backgrounds of much higher monovalent cation concentrations (6, 15-17)

While there are important examples of specifically bound ions that are required for RNA folding and function (6, 18), the vast majority of interacting ions are dynamically associated in a sheath that surrounds these molecules, referred as the “ion atmosphere” (7, 19-23). Unlike specifically bound ions that can be investigated by x-ray crystallography and other static structural techniques (18, 24-27), the dynamic ions present in the ion atmosphere are refractory to most traditional experimental methods (7, 20, 28). Yet, the ion atmosphere is a critical structural, dynamic, and energetic component of nucleic acids that profoundly affects their folding, compaction, and interactions. Hence, understanding RNA structure and function requires understanding the properties and energetics of its ion atmosphere.

An experimental approach that has been successful for studying the ion atmosphere around DNA and testing theoretical predictions is “ion counting” (20-23, 29, 30). It quantifies the number of thermodynamically accumulated cations and thermodynamically excluded anions around a negatively-charged macromolecule such as DNA. Particularly effective is ion counting through <u>b</u>uffer-<u>e</u>xchange combined with inductively <u>c</u>oupled <u>p</u>lasma <u>m</u>ass <u>s</u>pectroscopy (BE-ICPMS), as it allows the study of a large variety of ions over a broad range of ion concentrations, from tens of micromolar to molar (20, 21, 30). Prior studies have shown a strong preferential attraction of cations over the exclusion of anions—24-bp DNA attracted 37 ± 1 cations and excluded 9 ± 1 anions at 10 mM salt concentration, corresponding to 0.804 ± 0.02 attracted cation and 0.195 ± 0.02 excluded anion per charge unit of the double stranded (ds)DNA (20, 21, 29). Monovalent cation occupancy in the ion atmosphere is insensitive to the cation size across the alkali metal ions Na^+^, K^+^, Rb^+^, and Cs^+^, contrary to several computational predictions (30-33). Ion counting also revealed preferential association of divalent cations over monovalent cations around the dsDNA; e.g., with Na^+^ in 4-fold excess of Mg^2+^ (20 vs 6 mM), the ion atmosphere nevertheless has 4-fold more Mg^2+^ than Na^+^ (20, 30).

Over the past decades, experimental and computational studies have considerably advanced our understanding of the ion atmosphere around DNA duplexes. Yet, our knowledge of the ion atmosphere around RNA helices is limited. There are several computational studies dedicated to quantifying the RNA-ion interactions within the ion atmosphere (34-40); in particular, Poisson-Boltzmann (PB) calculations have emerged as the approach of choice, in part because it is easily implementable, computationally tractable, and conceptually straightforward (41-46). However, there are few experimental studies on the RNA electrostatics, and the complex RNAs used typically prevents isolating and dissecting the ion atmosphere and its associated energetics (41-48). Prior theoretical and computational studies have highlighted a higher linear charge density of the dsRNA compared to the dsDNA that is predicted to result in a stronger electrostatic field around dsRNAs (49-52) and stronger with ions, and in particular with divalent cations like Mg^2+^ (34-36, 39, 42).

Given the general important of RNA in biology and the motivation to better understand its electrostatic properties, we carried out ion counting experiments for monovalent and divalent cations around a 24-bp RNA. We compared its ion atmosphere composition with our previous results for a 24-bp DNA having the same sequence composition. We also compared the experimental results to theoretical Poisson-Boltzmann (PB) predictions of the ions within the RNA’s ion atmosphere. Our ion counting results support the predicted stronger electrostatic field of dsRNA than dsDNA and, as was previously observed for dsDNA, results for monovalent cations agree with PB predictions whereas those for Mg^2+^ do not.

## MATERIAL AND METHODS

### Reagents

DNA and RNA oligonucleotides were purchased from IDT (Integrated DNA Technologies, USA). The following DNA sequences were used <u>24S1</u>: _5′_GGT GAC GAG TGA GCT ACT GGG CGG_3′_, <u>24S2</u>: _5′_CCG CCC AGT AGC TCA CTC GTC ACC_3′_, 23S1: _5′_GGT GAC GAG TGA GCT ACT GGG CG_3′_ and 23S2: _5′_CGC CCA GTA GCT CAC TCG TCA CC_3′_. 24mer RNA sequences were the same as 24S1 and 24S2 except for containing uracil instead of thymine bases. All salts were of the highest purity (TraceSELECT® or BioXtra, Sigma-Aldrich USA). All solutions were prepared in high purity water, ultra-low TOC biological grade (Aqua Solutions, USA).

### Preparation of DNA and RNA samples

DNA and RNA constructs used in this study were duplexes assembled from chemically synthesized oligonucleotides. Prior to assembly, oligonucleotides were purified by reverse-phase HPLC (XBridge Oligonucleotide BEH C18; Waters, MA) and desalted using centrifugal Amicon Ultra-3K filters. The DNA and RNA constructs were prepared as described previously(20, 21).

### Buffer Equilibration-Inductively Coupled Plasma Mass Spectroscopy (BE-ICPMS)

Buffer equilibration for DNA and RNA was carried out using Amicon Ultracel-30K filters (Millipore, MA). Salt samples were prepared in 2 mM Na-EPPS or Mg-EPPS, pH 8.5 and their concentrations were determined by ICP MS. The initial 500 μL of 0.2 to 2 mM DNA or RNA samples, with the salt of interest, was spun down to ∼100 μL at 7000 x g in Amicon Ultracel-30K filters at 4 °C (to minimize solution evaporation) (53). As shown previously, equilibration between ions associated with nucleic acids and the bulk ions was completed after five rounds of the buffer exchange without any loss of the DNA or RNA; no DNA or RNA was detected in flow-through samples, as determined by ICP MS (21).

### Ion counting

Inductively coupled plasma mass spectrometry (ICPMS) measurements were carried out using a XSERIES 2 ICPMS (Thermo Scientific, USA). Samples were analyzed as described in references (20, 21, 53). Briefly, aliquots (5–20 μL) of DNA- or RNA-containing sample, the flow-through from the final equilibration, and the equilibration buffer were diluted to 5 mL in 15 mL Falcon tubes with water. Dilution factors, the ratio of diluted to total sample volume, were used to maintain sample concentrations within the linear dynamic range of detection. Calibrations were carried out using standards from SpexCertiPrep (USA). Quality control samples, containing each element of interest at 50 μM, were assayed every ten samples to estimate measurement precision (21, 53). A solution of 5% ammonium hydroxide in highly pure, ion-free water (Mili Q) was used as a wash-out solution between measurements (54).

Ion counting data point reported were collected from 2-3 independent experiments (i.e. ‘biological’ replicate). Errors are the standard deviation of all biological and technical replicates for a given sample.

The number of associated ions around the DNA- and RNA-duplex is reported here as a preferential interaction coefficient Γ_i_ (i = + or □, indicating cation or anion, respectively), where Γ_i_ is the difference in the ion concentration between the equilibrated nucleic acid-containing sample 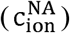 and the bulk solution 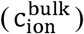, divided by the DNA or RNA concentration (c_NA_; determined by phosphorous measurements using ICPMS) (eq 1).

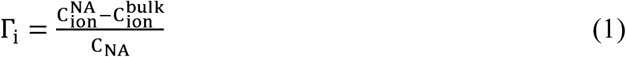

For DNA or RNA, the cation preferential interaction coefficient, Γ_+_, is expected to be greater than zero, indicating their accumulation around the negatively charged polyelectrolytes, and Γ_−_ for an anion is expected to be less than zero due to repulsive interactions with the DNA or RNA.

### Quantification of cation competition

To evaluate differences in the association between Mg^2+^ and monovalent cations (M^+^) with 24-bp RNA we used the same method as described previously (20). Subsequently, we compared these results to experimental data for the Mg^2+^ vs. M^+^ competition around a 24-bp DNA, from the same method and published previously.^30^ Briefly, the number of competing cations (CC) and Mg^2+^ cations around the RNA was measured over a range of CC concentrations at a fixed Mg^2+^ concentration of 6 mM. The competition constant ([CC]_1/2_) was defined as the concentration of competing cation at which the number of the CC cation and Mg^2+^ ions within the ion atmosphere are equal.

### Poisson Boltzmann (PB) calculations

The B-form 24-bp DNA and 23-bp DNA and A-form of 24-bp RNA were constructed with the Nucleic Acid Builder (NAB) package (55). Charges were assigned using the PDB2PQR routine (56) with the CHARMM parameter set. PB calculations were carried out using the Adaptive Poisson-Boltzmann Solver (APBS, version 1.4.1) (57) on a 405 × 405 × 578 Å^3^ grid with a grid spacing of 1.8 Å and the ion size equal 2 Å. As ion counting experiments were carried out at 4 °C, the simulation temperature was set to 277.15 K and the dielectric constant of the solvent was set to 86, characteristic of water at 4 °C (58). The internal dielectric of the DNA and the RNA was set to 2. The solvent-excluded volume of the DNA and the RNA molecules was defined with a solvent probe radius of 1.4 Å. Boundary conditions were obtained by Debye-Hückel approximation.

The preferential interaction coefficient of ions *i* of valence *z*_*i*_ associated with the DNA and the RNA was computed by integrating the excess ion density (3, 20, 59):

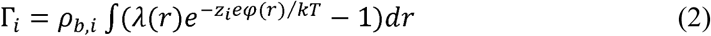

where *ρ*_*b,i*_ is the bulk ion density, *λ*(*r*) is an accessibility factor that defines the region in space that are accessible to ions (where *λ*(*r*) = 1 and *λ*(*r*) = 0 for the solvent-excluded region –i.e., inside the macromolecule), *e* is the elementary charge, *φ*(*r*) is the electrostatic potential, *k* is the Boltzmann constant, and *T* is the temperature.

The integration volume was defined as the entire volume of a simulation box including the solvent-excluded region in the DNA and the RNA interior (60). This approach matches the conditions for the experimental measurement, as the experiments employ equal total volumes for the nucleic acids and bulk reference samples.

## RESULTS

### RNA duplex accumulates more Na^+^ ions than a DNA duplex

To determine and compare electrostatic properties of DNA and RNA duplexes, we quantified the composition of ion atmospheres around the molecules by carrying out ion counting experiments for NaBr. We chose NaBr for its accuracy of detection by mass spectrometry and because it behaves similarly to the more physiological K^+^ and Cl^−^ ions (21, 30). Measurements revealed that 24-bp RNA attracts on average 2 more Na^+^ cations and excludes 2 less Br^−^ anions than dsDNA in the concentration rage of 10-500 mM (Figure 1A and Table S1 in Supplementary Information), despite the same overall charge of –46e and the same sequence composition. Under all experimental conditions, the sum of ionic charges (e.g. Na^+^ and Br^−^) from the ion atmosphere agrees well with the overall charge of 24-bp DNA and 24-bp RNA (Γ = +46; squares vs dashed lines in Figure 1A and Table S1 in Supplementary Information), as expected from the charge neutrality principle (20, 21).

**Figure 1.**
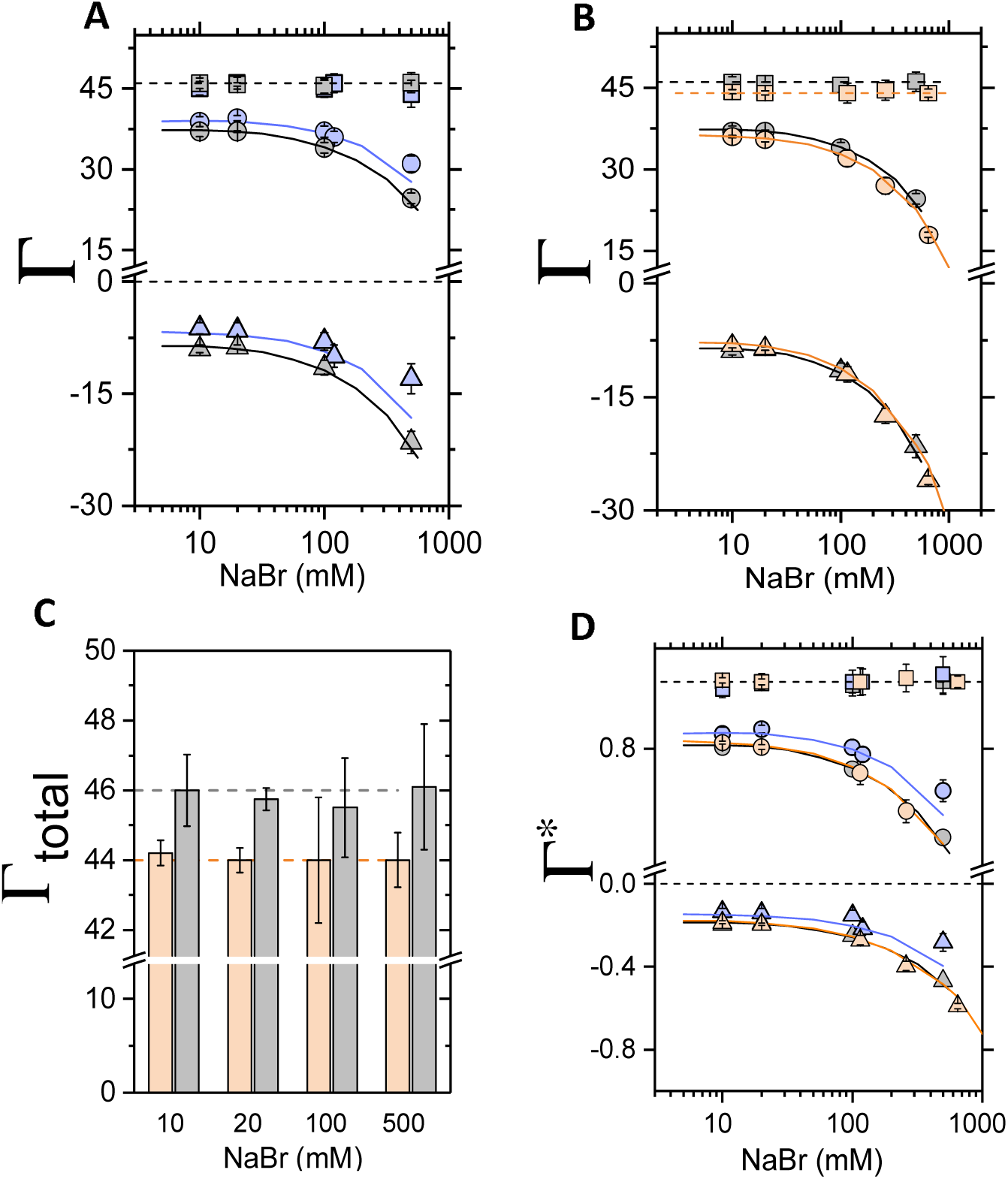
Quantification of the ion atmosphere around 24-bp RNA, 24-bp DNA, and 23-bp DNA duplexes. A) The preferential interaction coefficient (Γ) of attracted Na^+^ cations and excluded Br^−^ anions around 24-bp RNA (blue symbols) and 24-bp DNA (grey symbols). B) The preferential interaction coefficient of attracted Na^+^ cations and excluded Br^−^ anions around 23-bp (orange symbols) and 24-bp DNA (grey symbols). In (A) and (B), the total charge of the ion atmosphere summed from the individual ion measurements is shown as squares, and the dashed lines at Γ = +46 and Γ = +44 represent the theoretical charge needed to neutralize the 24-bp DNA and 24-bp RNA charge of −46e and 23-bp DNA charge of −44e. C) The total number of ions within the ion atmosphere around 23-bp (orange bars) and 24-bp DNA (grey bars) from (B). Dashed lines as in (B). D) The fraction of attracted Na^+^ cations and excluded Br^−^ anions per negative charge (phosphorous group) of the molecule, as defined by equation 3. Experimental results from BE-ICPMS ion counting are compared to PB predictions for 24-bp dsRNA (—), dsDNA: 24-bp (—) and 23-bp (—). Each data point is the average of two repeats from at least two independent experiments. The reported errors are the standard deviations of all biological and technical replicates for a given sample. See Table S1-S3 in Supplementary Information for data.

The difference in the number of ions in the ion atmosphere around the RNA and DNA is relatively small, yet the two-samples t-test reveals that all data points are significantly different with p value < 0.02 except the data from 10 mM (p = 0.08). To further test that ion counting method can detect differences on this scale, we carried out the analogous experiments with 23-bp DNA (Figure 1B). The theoretical charge of the 23-bp DNA is –44e (e.g. 2e charge less than the 24-bp DNA) and the experimentally determined charge agreed well with this value (Γ = +44; orange squares vs the orange dashed line in Figure 1B and 1C). We also measured less Na^+^ cations attracted to and fewer Br^−^ excluded from the 23-bp DNA (t-test: p < 0.03 expect the data from 10 mM, p = 0.1). These results indicate that ion counting can resolve differences in the molecule charge as small as 2e. Further, results for NaBr association around 24-bp DNA present herein are in excellent agreement with previously published data, supporting the robustness of the BE-ICP MS method (Figure S1 and Tables S1 and S2) (21,30,64)

To compare electrostatic properties of RNA and DNA we represent the fraction of charge neutralization form associated cations 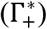 and from excluded anions 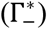 per unit charge, eq.

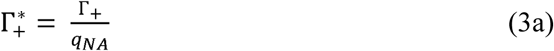

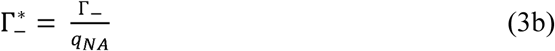

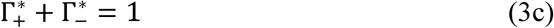

where, q_NA_ is the total charge of dsDNA or dsRNA, Γ_+_, and Γ_−_ are preferential coefficients for cations and anions respectively, as defined above. The sum of 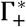 and 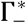 must equal one, following the charge neutrality principle.

We observed that the fraction of the associated Na^+^ around 24-bp RNA is larger than for 24-bp DNA (Figure 1D): 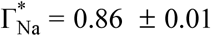 vs. 0.80 ± 0.004 for RNA and DNA, respectively (20 mM NaBr). In contrast we measured no difference in the association of Na^+^ between 24-bp and 23-bp DNA: 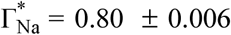 vs. 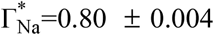 for 23-bp and 24-bp DNA, respectively (20 mM NaBr). A greater attraction of cations and the weaker repulsion of anions is indicative of a stronger electrostatic potential around dsRNA than around dsDNA (49-52), and this stronger electrostatic potential is predicted theoretically by the Poisson-Boltzmann (PB) equation (20, 60-62). Indeed, there is excellent quantitative agreement between the experimental data and the Poisson-Boltzmann predictions (Figure 1A, 1C, and 1D, points vs. lines).

Counting ions around charged molecules can also be achieved by anomalous X-ray scattering (ASAXS) (22, 29, 41). Our ion counting data give results similar to previous ASAXS data for monovalent ions around 25-bp DNA and 25-bp RNA (29, 34) but provide higher precision (Figure S2 and Table S3 in Supplementary Information). These differences could be experimental or result from sequence differences. We note that the high-throughput and explicit counting of ions via BE-ICPMS likely renders it a preferable tool for broad investigation of ion atmosphere contents as well as for carrying out rigorous error estimates; it is also more directly assays ion numbers, but it does not provide information about ion distributions that can be obtained from ASAXS (7, 29, 63).

### RNA duplex interacts significantly stronger with Mg^2+^ compared to a DNA duplex

To provide an independent test for the electrostatic differences between the dsDNA and dsRNA we measured monovalent cation competition for binding to dsDNA and dsRNA against constant concentration of Mg^2+^. DNA and RNA preferentially interact with divalent cations [M^2+^] over monovalent cation [M^+^], and the preference for divalent over monovalent increases as the strength of the molecule’s electrostatic field increases (15, 35, 36, 39, 41, 64, 65).

Our previous ion counting measurements of Mg^2+^ association with the dsDNA revealed 21.5 ± 0.5 divalent cations around the molecule for solutions containing only Mg^2+^ cations (20, 30). We measured the similar number of attracted Mg^2+^ around 24-bp RNA, Γ_Mg_ = 22.0 ± 0.5. We carried out [Mg^2+^] vs. [M^+^] competition experiments against two monovalent cations Na^+^ and Cs^+^ as competing cations [CC] to address the effect of the [CC] size (e.g. Cs^+^ is larger than Na^+^ by comparing their ionic radius) in addition to the charge (e.g. monovalent vs. divalent) on energetics of the cation association. Upon increasing the bulk concentration of Na^+^ or Cs^+^ the number of associated Mg^2+^ decreased and the number of the competing monovalent cations increased. As expected, charge neutrality was maintained across all concentrations (Figure 2A and 2B). To estimate energetics of monovalent cation interacts with the DNA or the RNA, relative to a Mg^2+^ background, we introduce the unitless parameter α, defined as:

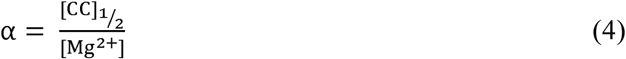

where [CC]_1/2_ is the cation competition constant—i.e., the concentration of competing cation (CC) at which the number of the CC cation and Mg^2+^ ions within the ion atmosphere are equal, and [Mg^2+^] is the concentration of the background Mg^2+^, here 6 mM. The relative preferential cation occupancy from the data of Figures 2A and 2B and Table S4 is summarized in Figure 2C in terms of α. The cation competition experiments show that Mg^2+^ interacts stronger with the 24-bp RNA compared to the 24-bp DNA—with a measured α value that is 2-fold higher for the RNA (Figure 2C and Table S4). Similar values of α for Na^+^ and Cs^+^ against Mg^2+^ suggest that the occupancy of the monovalent cations within the ion atmosphere is insensitive to their size, as shown previously in M^+^ vs. M^+^ cation competition experiments.^30^

**Figure 2.**
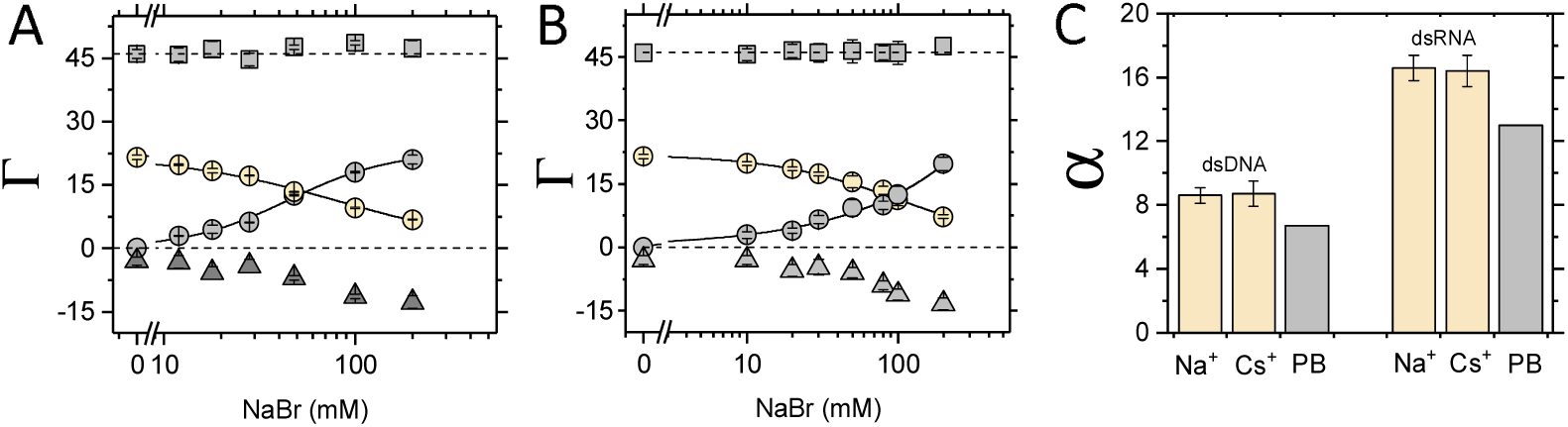
Competitive association of monovalent cations against Mg^2+^ for a 24-bp DNA and RNA duplexes. (A,B) The preferential interaction coefficient (Γ) of associated Na^+^ cations (gray circles), replaced Mg^2+^ cations (yellow circles, [MgBr_2_] = 6 mM) and excluded Br^−^ anions (grey triangles) around the 24-bp DNA (A) and around the 24-bp RNA (B). In (A) and (B), the total charge of the ion atmosphere summed from the individual ion measurements is shown as squares, and the dashed lines at Γ = +46 represent the charge needed to neutralize the total 24-bp DNA and 24-bp RNA charge of −46e. Solid lines are fits to the Hill equation to provide an empirical guide. (C) Cation competition constants (*α*) for monovalent cations against 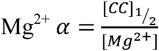 (eq. 4), where [CC]_1/2_ is the concentration of competing cation at which the number of the CC cation and Mg^2+^ ions within the ion atmosphere are equal, [Mg^2+^] is the background concentration, here [Mg^2+^] = 6 mM. Each data point in (B-C) with the 24-bp RNA is the average of three independent measurements. Error bars as in Figure 1. See Table S4 in Supplementary Information for data. Ion counting data for the 24-bp DNA (A and C) are from reference (30), where the same methodology was used.

We also carried out PB calculations for the divalent vs. monovalent cation competition binding (Figure 2C). PB predicts that dsRNA interacts stronger with Mg^2+^ than dsDNA (e.g. *α*_*DNA*_ 6.7 and *α*_*RNA*_ 13.6 in Figure 2C) and that twice as much monovalent ion concentration as for the dsDNA is required to reach the equal amount of divalent and monovalent cations within the ion atmosphere around the dsRNA, in accord with the experimental results (Figure 1C). However, PB underestimates the strength of the Mg^2+^-DNA and Mg^2+^-RNA interactions, predicting lower monovalent cation concentrations are required to replace Mg^2+^ than is observed experimentally (Figure 1C).

In summary, experimental data show that dsRNA interacts stronger with Mg^2+^ than dsDNA, consistent with theoretical predictions, but there are nevertheless quantitative differences between experiment and theoretical results.

## DISCUSSION

Our ion counting experimental studies allow us to evaluate electrostatic properties of RNA and DNA duplexes. We observed that dsRNA attracts more monovalent cations than dsDNA and interacts more favourably with Mg^2+^. These results indicate that the electrostatic field around the RNA duplex is stronger than that of the DNA duplex, as proposed by numerous computational studies (34-39, 65).

To illustrate the electrostatic differences between RNA and DNA, we carried out PB calculations of the electrostatic surface potentials for the respective canonical helices (Figure 3B). PB predicts the electrostatic potential at the phosphate backbone is higher for the dsRNA than for the dsDNA, –830 mV vs. –640 mV (at 10 mM NaCl). This difference is visualized in Figure 3B, where the deeper red colour indicates the larger negative electrostatic potential around the phosphate groups of the dsRNA.

**Figure 3.**
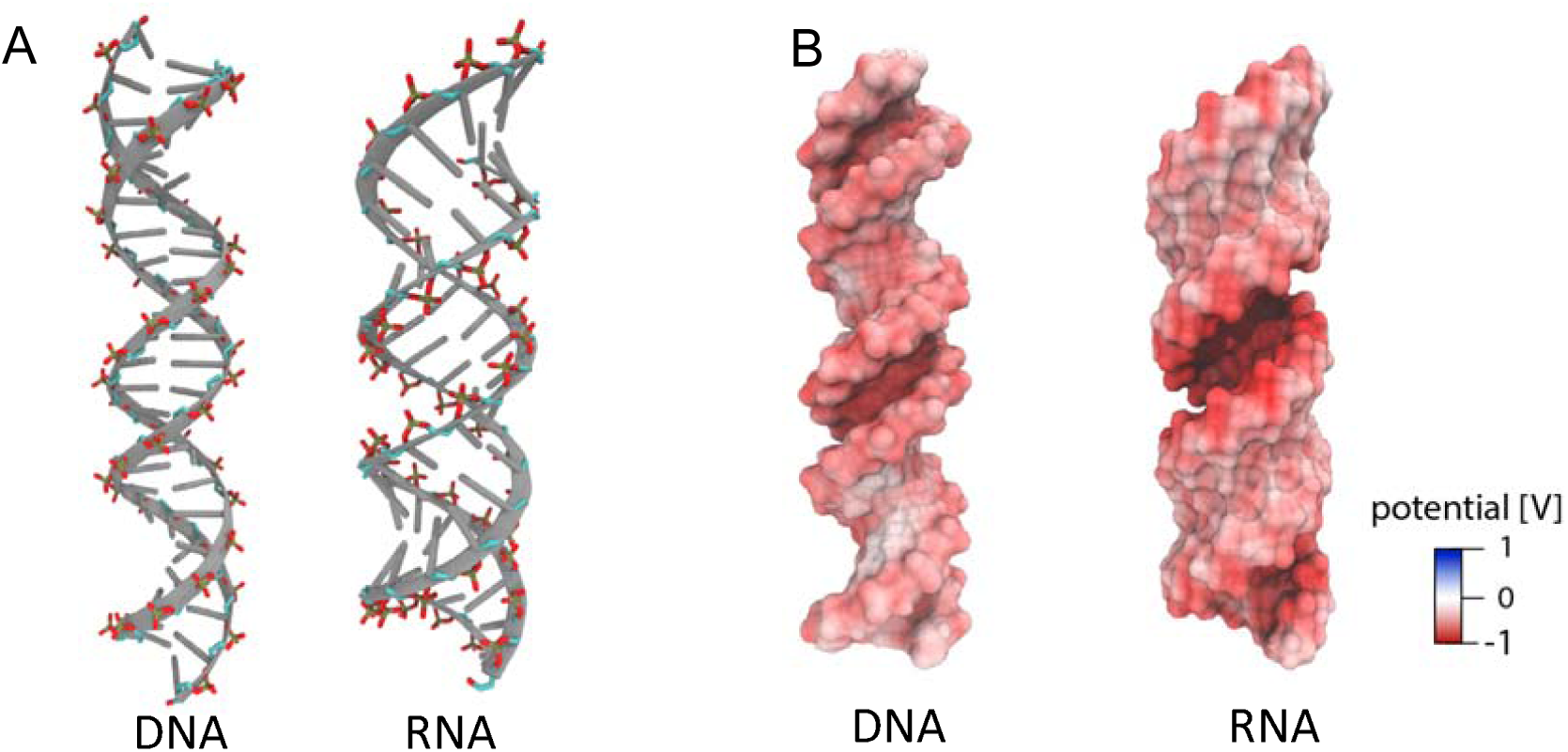
Comparison of DNA and RNA duplex shape and electrostatic potential. A) DNA and RNA duplexes with highlighted phosphate residues. The B-form 24-bp DNA duplex and A-form of 24-bp RNA were constructed with the Nucleic Acid Builder (NAB) package (55). B) Poisson-Boltzmann calculations of electrostatic surface potential of the DNA and RNA duplexes from (A). The electrostatic potential mapped to the molecular surfaces was calculated using Adaptive Poisson-Boltzmann Solver (APBS) (56) and the figures were rendered with Visual Molecular Dynamics (VMD) (66).

Topological differences between DNA and RNA duplexes play the major role in defining the electrostatic properties of the molecules. Phosphoryl groups in dsRNA face inwards, along a surface plane of the major groove, whereas those in dsDNA are oriented toward bulk solvent (Figure 3A). The shorter P–P distances along dsRNA (5.65 Å vs 6.62 Å for dsRNA and dsDNA, respectively) and across (9.97 Å vs 11.69 Å for dsRNA and dsDNA, respectively) the major groove for dsRNA and the minor groove for dsDNA result in the stronger RNA electrostatic field (Figure 3B), as Coulombic interactions are distance dependent. The inward-facing of the negatively charged phosphate groups may also contribute to enhancing of the electrostatic potential through focusing electrostatic field lines, a phenomenon referred as electrostatic focusing (67). Indeed, previous crystallographic studies have shown stronger localization of the Mg^2+^ within the deep major groove and at the phosphate groups bridging across the other month of the narrow major groove, at the most negative electrostatic potential regions of A-form duplex (65).

An important step towards developing quantitative and predictive models of RNA structure, folding and interactions with binding partners is the quantitative understanding of nucleic acid–ion interaction within the ion atmosphere and at the specific ion binding site. Given the highly complexity and dynamic nature of the ion atmosphere, understanding will require synergy between theory and experiment. Experimental methods like ion counting can quantify the overall content of the ion atmosphere and energetics of competitive association of cations but does not provide information about the distribution of ions within the atmosphere. Computational models can in principle provide a thorough and deep understanding of ion-nucleic acid interactions, solvent-nucleic acid interactions, and the dynamic and energetic consequences of these interactions (31, 35, 60, 68-75). However, the accuracy of such models cannot be assumed, and even matching to one a few prior experimental measurements in insufficient to establish the veracity of models for systems as complex and multi-variant as nucleic acid/ion interactions in solution. Instead, robust and deep tests of *bona fide* blind predictions are needed, via ion counting and additional experimental methods.

Previous studies on competitive association of monovalent cations with the dsDNA have proven invaluable in testing computational models (30). E.g., monovalent cation occupancy in the dsDNA and dsRNA ion atmosphere is insensitive to the cation size across the alkali metal ions Na^+^, K^+^, Rb^+^, and Cs^+^, contrary to all-atom computational models and highlighting the need to reevaluate molecular mechanical force fields for solute–solvent and solvent– solvent interactions. Our new experimental results for dsRNA-ion interactions provide the opportunity to test newly developed all-atom models of RNA-Mg^2+^ interactions (70, 72, 74, 76, 77) and initiate a feedback loop between computation and experiment.

## ACKNOWLEDGEMENT

We thank members of the Herschlag lab for helpful discussions and for critical advice.

