## Supplemental information for "Quantitative studies of an RNA duplex electrostatics by ion counting"

M. G. and D. H

**Overview of the Supplementary Information**

In this supplementary information, we provide a table with preferential ion interaction coefficients $\Gamma_{i}$ (e.g. the number of associated ions, i = Na^+^ or Br^−^), around 24-bp RNA, 24-bp and 23-bp DNA (Table S1) for NaBr; a table summarizing the fraction of charge neutralization from attraction of Na^+^ measured by ASAX and BE-ICPMS around dsRNA and dsDNA (Table S2); a table with preferential ion interaction coefficients from competition experiments between Na^+^:Mg^2+^ and Cs^+^:Mg^2+^ around 24-bp RNA. The results are consistent with observations in the main text and support the conclusions described therein.

**Table S1:** Experimentally determined preferential interaction coefficients ($\Gamma_{i}$) for NaBr around 24-bp RNA, 24-bp DNA, and 23-bp DNA

|  | **24 bp RNA** | | | **24 bp DNA** | | | **23 bp DNA** | | |
| --- | --- | --- | --- | --- | --- | --- | --- | --- | --- |
| **C [M]** | $\boldsymbol{\Gamma}_{\boldsymbol{Na}^{\boldsymbol{+}}}$ | $\boldsymbol{\Gamma}_{\boldsymbol{Br}^{\boldsymbol{-}}}$ | **total** | $\boldsymbol{\Gamma}_{\boldsymbol{Na}^{\boldsymbol{+}}}$ | $\boldsymbol{\Gamma}_{\boldsymbol{Br}^{\boldsymbol{-}}}$ | **total** | $\boldsymbol{\Gamma}_{\boldsymbol{Na}^{\boldsymbol{+}}}$ | $\boldsymbol{\Gamma}_{\boldsymbol{Br}^{\boldsymbol{-}}}$ | **total** |
| 0.01 | 39 ± 1.0 | -6.0 ± 0.7 | 45.0 ± 1.2 | 37.0 ± 1.0 | -9.0 ± 0.5 | 46.0 ± 1.0 | 36.0 ± 0.2 | -8.0 ± 0.3 | 44.0 ± 0.3 |
| 0.02 | 39.5 ± 0.5 | -6.5 ± 1.0 | 46.0 ± 1.0 | 37.0 ± 0.2 | -8.75 ± 0.2 | 46.0 ± 0.3 | 35.0 ± 0.3 | -8.6 ± 0.2 | 44.0 ± 0.3 |
| 0.10 | 37.0 ± 1.0 | -8.0 ± 1.2 | 45.0 ± 1.6 | 34.0 ± 1.0 | -11.5 ± 1.0 | 45.5 ± 1.4 | 32.0 ± 1.0 | -12.0 ± 1.0 | 44.0 ± 1.0 |
| 0.12 | 36.0 ± 1.0 | -10.0 ± 1.5 | 46.0 ± 1.8 | - | - | - |  |  |  |
| 0.26 | - | - | - | - | - | - | 27.0 ± 1.5 | -17.5 ± 1.0 | 44.5 ± 1.8 |
| 0.50 | 31.0 ± 1.5 | -13 ± 2.0 | 44.0 ± 2.5 | 24.6 ± 1.0 | -21.5 ± 1.5 | 46.0 ± 1.8 |  |  |  |
| 0.65 | - | - | - |  |  |  | 18.0 ± 0.5 | -26 ± 0.5 | 44.0 ± 0.5 |

**Table S2.** Interaction coefficients for NaBr around 24-bp DNA obtained previously in reference (3)

|  | **NaBr** | | |
| --- | --- | --- | --- |
| **C [M]** | $\boldsymbol{\Gamma}_{\boldsymbol{Na}^{\boldsymbol{+}}}$ | $\boldsymbol{\Gamma}_{\boldsymbol{Br}^{\boldsymbol{-}}}$ | **total** |
| 0.010 | 37.0 ± 0.9 | -9.0 ± 0.9 | 46 ± 1.3 |
| 0.050 | 36.0 ± 0.7 | -8.7 ± 0.7 | 44.7 ± 1.0 |
| 0.100 | 35.0 ± 1.0 | -10 ± 1.0 | 45 ± 1.4 |
| 0.200 | 32.0 ± 1.5 | -14.5 ± 1.5 | 46.5 ± 2.0 |
| 0.350 | 28.0 ± 1.5 | -16.7 ± 1.2 | 44.7 ± 2.0 |
| 0.500 | 24.6 ± 1.0 | -21.5 ± 1.5 | 46.1 ± 1.8 |

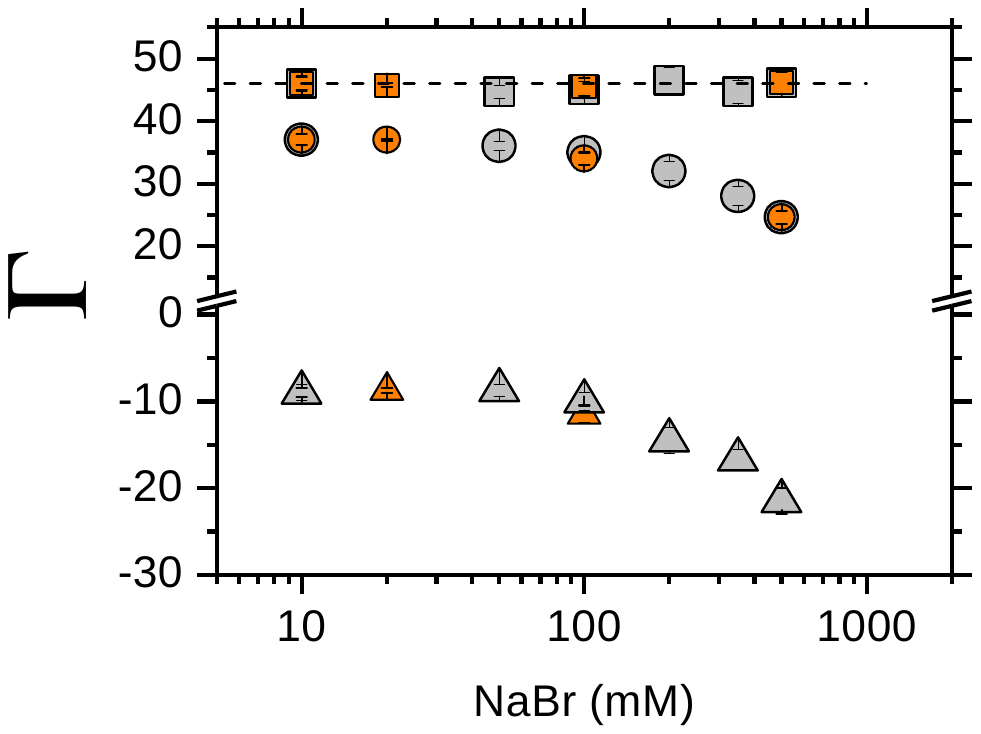

**Figure S1**. Comparison of current (orange symbols) and previous (grey symbols) ion counting results for association of NaBr around 24-bp DNA from BE-ICP MS measurements. Data point in grey are from reference (3) and values are given in Table S2.

**Table S3:** Experimentally determined fraction of charge neutralization ($\Gamma_{Na}^{*}$) for Na^+^ around dsRNA and dsDNA.

|  | **dsRNA** | | **ds DNA** | |
| --- | --- | --- | --- | --- |
| **C [M]** | ${\boldsymbol{\Gamma}^{\boldsymbol{*}}}_{\boldsymbol{Na}}^{\boldsymbol{ASAXS}}$ | ${\boldsymbol{\Gamma}^{\boldsymbol{*}}}_{\boldsymbol{Na}}^{\boldsymbol{BE-ICPMS}}$ | ${\boldsymbol{\Gamma}^{\boldsymbol{*}}}_{\boldsymbol{Na}}^{\boldsymbol{ASAXS}}$ | ${\boldsymbol{\Gamma}^{\boldsymbol{*}}}_{\boldsymbol{Na}}^{\boldsymbol{BE-ICPMS}}$ |
| 0.1 | 0.73 ± 0.06 ^(a)^ | 0.80 ± 0.02 | 0.71 ± 0.06 ^(b)^ | 0.74 ± 0.02 |

a) Data taken from reference 1

b) Data taken from reference 2

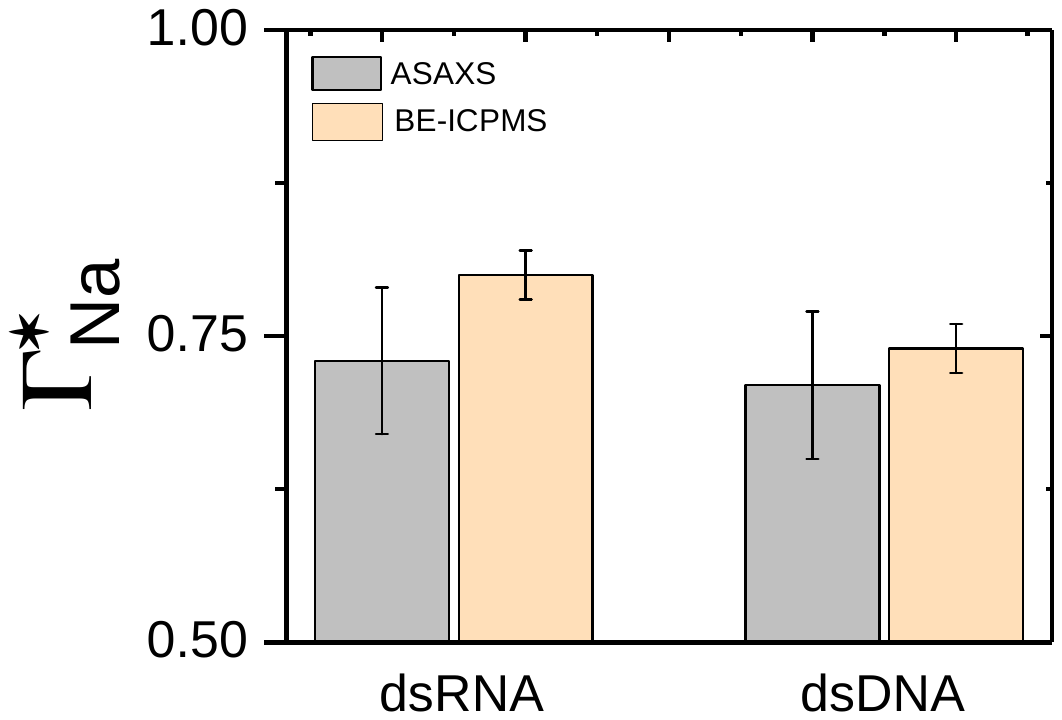

Figure S2. Comparison of experimentally determined fraction of charge neutralization ($\Gamma_{Na}^{*}$) for Na^+^ around dsRNA and dsDNA from ASAXS and BE-ICP MS. Data from Table S2.

**Table S4:** Experimentally determined preferential interaction coefficients and α value for **NaBr** and **CsBr** around 24-bp RNA in the presence of 6 mM MgBr_2_.

| **NaBr** | **24-bp RNA** | | | | **CsBr** | **24-bp RNA** | | | |
| --- | --- | --- | --- | --- | --- | --- | --- | --- | --- |
| **C [M]** | $\boldsymbol{\Gamma}_{\boldsymbol{Na}^{\boldsymbol{+}}}$ | $\boldsymbol{\Gamma}_{\boldsymbol{Mg}^{\boldsymbol{2+}}}$ | $\boldsymbol{\Gamma}_{\boldsymbol{Br}^{\boldsymbol{-}}}$ | **total** | **C [M]** | $\boldsymbol{\Gamma}_{\boldsymbol{Cs}^{\boldsymbol{+}}}$ | $\boldsymbol{\Gamma}_{\boldsymbol{Mg}^{\boldsymbol{2+}}}$ | $\boldsymbol{\Gamma}_{\boldsymbol{Br}^{\boldsymbol{-}}}$ | **total** |
| 0.00 | 0 | 21.0 ± 0.5 | -3.0 ± 1.0 | 46.0 ± 1.0 | 0 | 0 | 21.0 ± 0.5 | -4.0 | 46.0 ± 0.7 |
| 0.015 | 0.6 ± 0.5 | 22.0 ± 0.5 | -2.0 ± 0.5 | 46.8 ± 0.8 | 0.02 | 3.6 ± 1.0 | 19.0 ± 0.5 | -5.4 ± 1.0 | 47.0 ± 1.5 |
| 0.01 | 2.9 ± 0.8 | 20.0 ± 0.4 | -3.0 ± 1.6 | 45.6 ± 1.4 | 0.03 | 8.0 ± 1.0 | 16.0 ± 0.3 | -5.0 ± 0.6 | 45.6 ± 1.2 |
| 0.02 | 3.9 ± 0.6 | 18.5 ± 0.4 | -5.4 ± 1.4 | 46.5 ± 1.6 | 0.06 | 10.0 ± 0.5 | 16.0 ± 0.6 |  | 46.0 ± 1.2 |
| 0.03 | 6.5 ± 1.0 | 17.4 ± 0.4 | -4.7 ± 1.8 | 46.0 ± 2.0 | 0.11 | 11.5 ± 1.0 | 13.5 ± 0.8 | -4.0 ± 1.0 | 45.6 ± 1.6 |
| 0.05 | 9.3 ± 2.0 | 15.5 ± 1.4 | -6.0 ± 1.2 | 46.3 ± 2.7 | 0.2 | 19.0 ± 0.9 | 8.1 ± 1.4 | -10.0 ± 1.0 | 45.0 ± 1.7 |
| 0.08 | 10.0 ± 1.0 | 13.5 ± 1.0 | -9.0 ± 1.0 | 46.0 ± 1.7 |  | - | - | - | - |
| 0.10 | 12.5 ± 2.0 | 11.0 ± 1.3 | -11.0 ± 1.0 | 46.0 ± 2.7 |  | - | - | - | - |
| 0.20 | 19.8 ± 1.5 | 7.2 ± 0.4 | -13.3 ± 1.3 | 47.5 ± 2.0 |  | - | - | - | - |
|  | $\alpha_{Na}^{*}$ = = 16.6 ± 0.8 | | | |  | $\alpha_{Cs}^{*}$= 16.4 ± 1.0 | | | |

*Defined in the main text
